# Utility Analyses of AVITI Sequencing Chemistry

**DOI:** 10.1101/2024.04.18.590136

**Authors:** Silvia Liu, Caroline Obert, Yan-Ping Yu, Junhua Zhao, Bao-Guo Ren, Jia-Jun Liu, Kelly Wiseman, Benjamin J. Krajacich, Wenjia Wang, Kyle Metcalfe, Mat Smith, Tuval Ben-Yehezkel, Jian-Hua Luo

## Abstract

**Background:** DNA sequencing is a critical tool in modern biology. Over the last two decades, it has been revolutionized by the advent of massively parallel sequencing, leading to significant advances in the genome and transcriptome sequencing of various organisms. Nevertheless, challenges with accuracy, lack of competitive options and prohibitive costs associated with high throughput parallel short-read sequencing persist.

**Results:** Here, we conduct a comparative analysis using matched DNA and RNA short-reads assays between Element Biosciences’ AVITI and Illumina’s NextSeq 550 chemistries. Similar comparisons were evaluated for synthetic long-read sequencing for RNA and targeted single-cell transcripts between the AVITI and Illumina’s NovaSeq 6000. For both DNA and RNA short-read applications, the study found that the AVITI produced significantly higher per sequence quality scores. For PCR-free DNA libraries, we observed an average 89.7% lower experimentally determined error rate when using the AVITI chemistry, compared to the NextSeq 550. For short-read RNA quantification, AVITI platform had an average of 32.5% lower error rate than that for NextSeq 550. With regards to synthetic long-read mRNA and targeted synthetic long read single cell mRNA sequencing, both platforms’ respective chemistries performed comparably in quantification of genes and isoforms. The AVITI displayed a marginally lower error rate for long reads, with fewer chemistry-specific errors and a higher mutation detection rate.

**Conclusion:** These results point to the potential of the AVITI platform as a competitive candidate in high-throughput short read sequencing analyses when juxtaposed with the Illumina NextSeq 550.

## Introduction

DNA and RNA sequencing are essential applications for deeper understanding of modern biology. In the last 20 years, high-throughput massively parallel short-read sequencing platforms have formed the backbone in deciphering the genomes and transcriptomes of numerous organisms [1-3]. As short-read sequencing platforms entered the market, they offered an ever-evolving variety of sequencing mechanisms, throughput capacities, and accuracies [4-6]. From that plethora of options, Illumina emerged as the dominant manufacturer of short-read sequencing platforms. One of the most widely used sequencing platforms from the Illumina portfolio is the NextSeq. Although historically, there was a lack of an alternative platform which could offer similar capacity, versatility, and sequencing accuracy, 2022 saw the entry of several new short-read sequencer manufacturers to the market, including Singular Genomics, Ultima Genomics, and MGI [7].

Another recent entry to the short-read sequencing space was that by Element Biosciences, whose AVITI sequencer uses a novel technology known as avidity sequencing. Avidity sequencing utilizes a highly specific multivalent binding between fluorescence-labeled nucleotide polymers and DNA templates within the enzymatic pocket of the engineered DNA polymerase to identify each nucleotide in the DNA templates [8]. The high affinity binding between the nucleotide polymers and DNA templates lead to a reduction of the dissociation constant, a decrease in the amount of required sequencing reagents, and an increase in sequencing accuracy. However, the utility of such sequencing improvements remains to be established. In this report, we performed multiple control sample analyses using whole genome, transcriptome, synthetic long-read transcriptome, and targeted synthetic long-read single-cell transcriptome assays on both Illumina and Element Biosciences platforms. The results indicate that while comparable consensus-based long-read metrics were observed between NovaSeq 6000 and AVITI sequencing chemistries, avidity sequencing resulted in substantially better short-read sequencing quality and accuracy in comparison to that obtained when using Illumina’s NextSeq 550, while retaining similar quantification robustness.

## Results

The AVITI benchtop platform from Element Biosciences currently has a capacity of 1 billion polonies per flow cell. This output capacity is more than double that of Illumina’s similarly sized NextSeq 550 and NextSeq 1000, which have maximal outputs of 400 million clusters per run and is comparable to that of the newer released NextSeq 2000 which has a maximal output of 1.2 billion clusters. It should be noted, however, that the AVITI is a dual flow cell platform, so the total output is roughly double that of the NextSeq 2000. Although the latter two Illumina platforms were optimized for cluster brightness, reduction in channel cross talk, improved signal to noise ratios and data yield as compared to the NextSeq 550, all three platforms provide equivalent data quality with regards to key applications (https://www.illumina.com/content/dam/illumina-marketing/documents/products/appnotes/nextseq-1000-2000-data-concordance-app-note-970-2020-001.pdf). We utilized Illumina’s NextSeq 550 as the control platform for our short-read analyses, to investigate the utility of the AVITI chemistry, normalizing datasets sizes where appropriate. To achieve a robust comparison, we performed short-read sequencing to evaluate PCR-Free whole genome library preparations of *Escherichia coli* and mRNA library preparations of an ERCC standard for the evaluation of transcriptome sequencing.

To make direct comparisons of synthetic long-read sequencing from an ERCC standard and single-cell samples from a liver cancer patient (Figure 1), samples starting from the same initial library preparations were utilized in Illumina and Element Biosciences platform specific sequencing. As a NextSeq 2000 was not available for our evaluations, to generate a comparable number of short reads from a single sequencing run, we opted to assess the data from a single AVITI flow cell to that generated by a single lane on a S4 NovaSeq 6000 flow cell. The NovaSeq 6000 is a dual flow cell, production-scale platform, which has a maximum output of 10 billion clusters (or roughly 2.5 billion clusters per lane) per S4 flow cell. Although the platform is significantly larger than the various NextSeq models, similar base quality metrics have been observed (https://q.omnomics.com/ords/f?p=118:34:::::P34_S1,P34_S2:NextSeq%20500,NovaSeq%206000; https://q.omnomics.com/ords/f?p=118:34:::::P34_S1,P34_S2:NextSeq%202000,NovaSeq%206000); [9]. It should be noted, though, that the percent error rate is lower on the NovaSeq 6000 than the NextSeq 550 [10].

**Figure 1.**
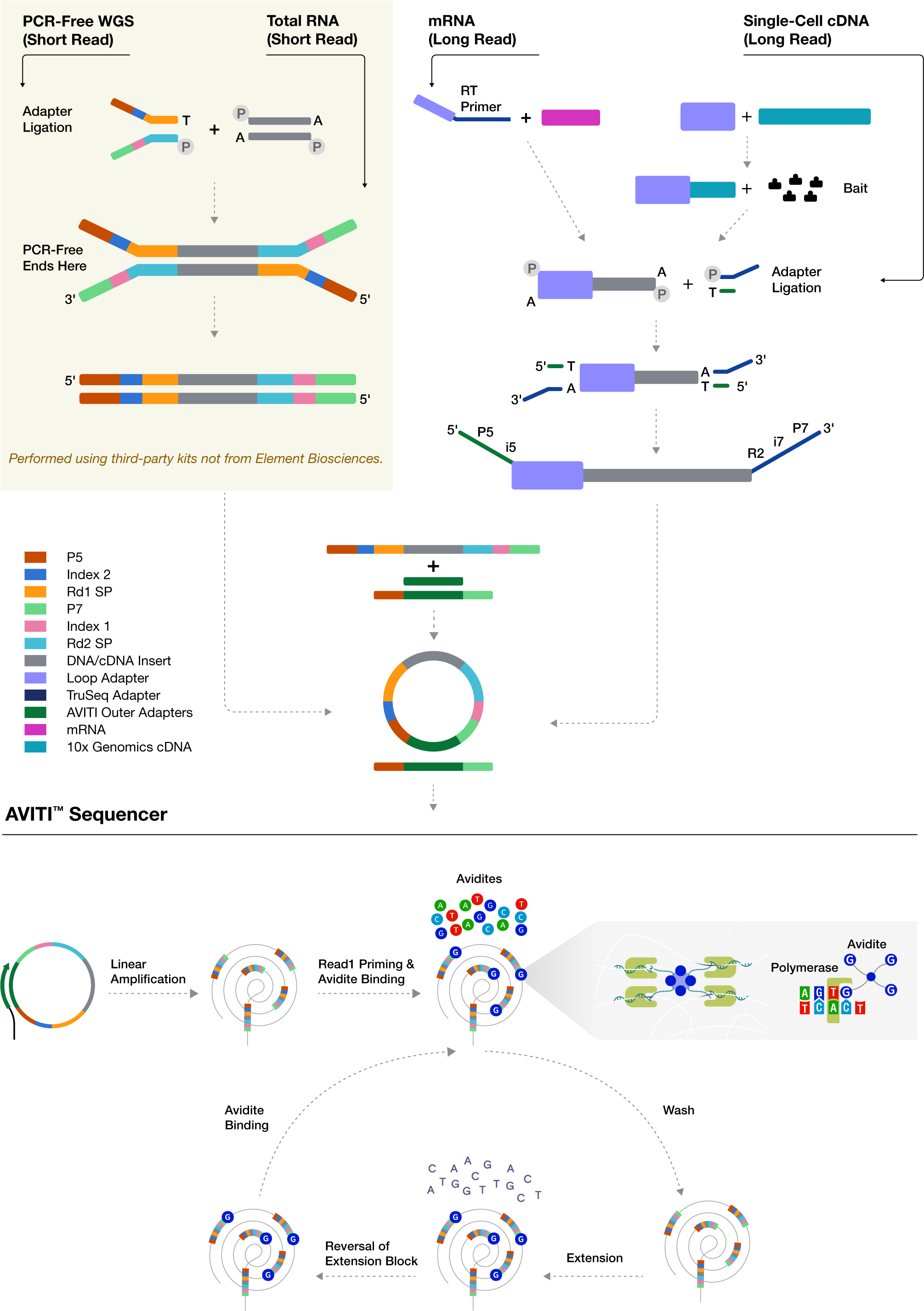
Schema of AVITI sequencing flowchart. PCR-free whole genome sequencing (WGS), transcriptome sequencing, long-read transcriptome sequencing, and targeted single-cell long-read sequencing processes are outlined. After linear amplification and circularization of the single stranded sequencing template, the process enters the cycle of sequencing process: avidite binding, wash, one base nucleotide extension, reversal of extension block.

### Short-Read Whole Genome Sequencing of the **E. coli** Genome

In this study, conducted from 2022 to 2024, 17 PCR-free libraries of the genome of *E.coli* strain B [CIP 103914, NCIB 11595, NRC 745] (∼4.6 Mb) were sequenced and analyzed with 2x150 cycles. The sequencing outputs from these runs on AVITI and NextSeq 550 were 3,429,918,215 and 2,204,183,238 paired end reads, respectively. As shown in Figure 2A, using avidity sequencing, over 91% of the reads from AVITI in read 1 obtained a per sequence quality score of 40 (equivalent to 0.01% error rate) using FASTQC [11]. The number of AVITI reads with quality score of 40 or higher dropped to 84.6% in read 2, suggesting that the majority of the reads had an accuracy of greater than 99.99%. As the NextSeq platforms only evaluate per base quality scores to Q30, no reads from the NextSeq 550 displayed a similar level of per sequence quality to that of the AVITI. When comparing the number of reads from both platforms that were able to achieve 99.9% accuracy (equivalent to per sequence quality scores of 30), the AVITI was able to ascribe 99.3% reads from read 1 and 97.3% reads from read 2. These results were statistically favorable in comparison with 93.5% reads from read 1 and 90.9% reads from read 2 off the NextSeq 550 (Supplemental Table 1).

**Figure 2.**
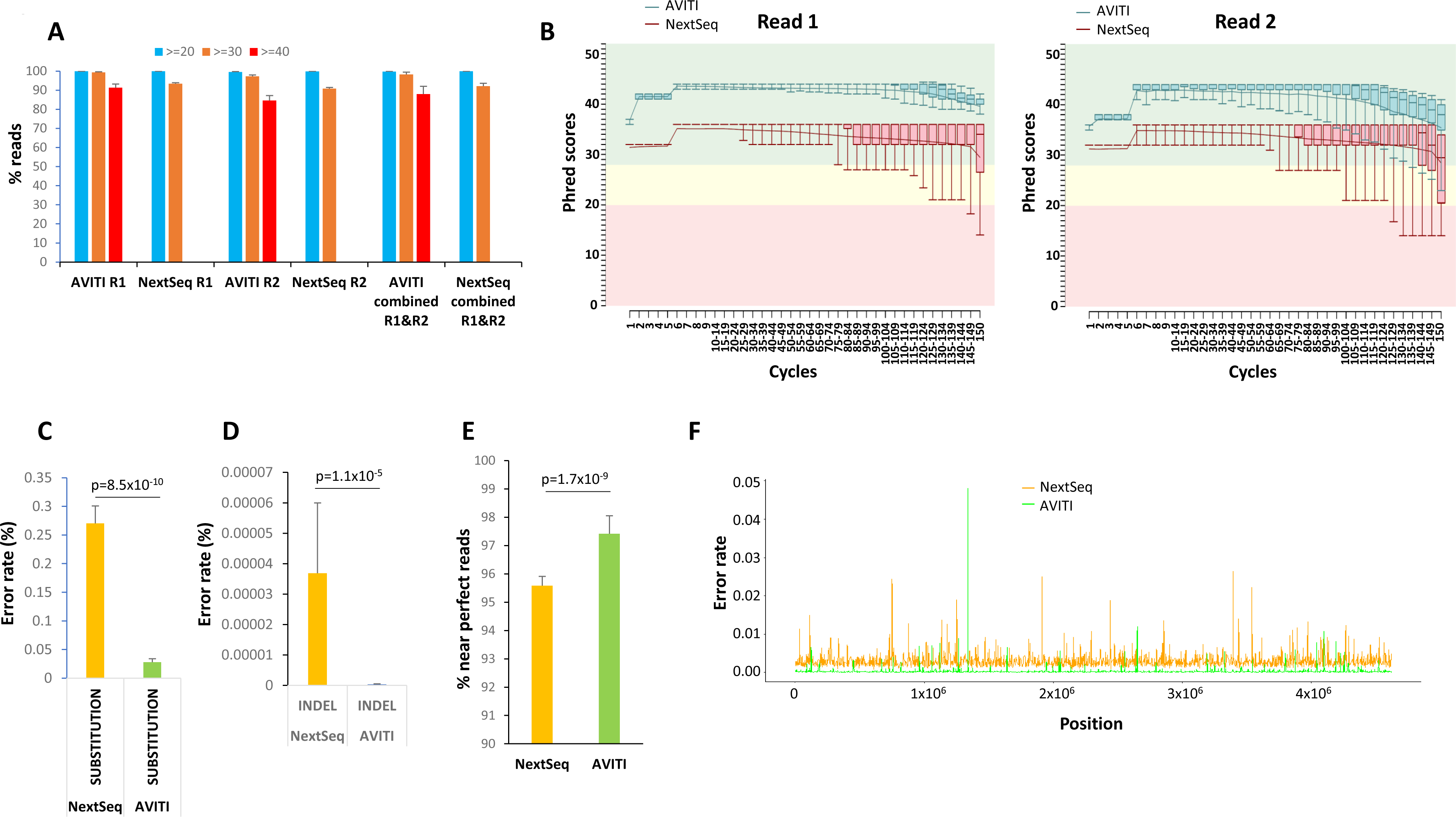
Sequencing of E. coli Genome by AVITI and NextSeq 550. (A) Average sequence quality score distributions of 17 samples of *E. coli* genome sequencing through AVITI and NextSeq 550 sequencing. Percentage of reads with scores more than 40, 30, and 20 scores are indicated. Standard deviations are shown. (B) Phred score distribution cycle by cycle of AVITI and NextSeq 550 of a matched sample sequencing of E. coli genome. Whisker box plots represent 10, 25, 50, 75 and 90 percentile distributions of reads in the indicated cycles. Both read 1 and read 2 were analyzed. (C) Percent substitution error rate of the mapped reads of matched 17 samples sequenced from AVITI and NextSeq 550. Standard deviations are shown. Wilcox p-value is indicated. (D) Percent insertion/deletion error rate of the mapped reads of matched 17 samples sequenced from AVITI and NextSeq 550. Standard deviations are shown. Wilcox p-value is indicated. (E) Percentage of the perfect mapped reads from AVITI and NextSeq 550. Standard deviations are shown. Wilcox p-value is indicated. (F) Histogram of 2000 loci error rate in the *E. coli* genome by AVITI and NextSeq 550.

To examine whether the quality differences between the two platforms were related to time-dependent decay in read quality, the Q scores were analyzed cycle by cycle. As shown in Figure 2B and Supplemental Figure 1, most of the median base Q scores in read 1 of AVITI sequences hovered above 40. The trend of the Q score started to drop after 100 cycles, with the lowest Q score slightly above 37 (99.98% accuracy) at the 150^th^ cycle. On the other hand, most of the average Q scores in read 1 of the NextSeq 550 sequences hovered in mid 30s with a wider range of quality variation. The trend line of Q scores on the NextSeq 550 began to drop after 80 cycles. A significant number of reads fell into abnormal (yellow) range after 120 cycles. Interestingly, there was no overlap of percentile read quality between the two platforms in read 1. For read 2, the Q scores were generally lower for both platforms. The Q scores from AVITI platform appeared to maintain in the normal range (green) for 134 cycles, while some bases fell into abnormal range after 135 cycles. On the other hand, the Q scores from the NextSeq 550 platform only maintained most of the reads in normal range up to 64 cycles. The downward trend started at 65 cycles. A significant number of base Q scores fell into abnormal and bad read (red) ranges after 65 cycles. A consistent big drop in Phred score occurred in the last two cycles of the sequencing for all the samples (Figure 2B and Supplemental Figure 1). Thus, there was a significant quality gap between the chemistries from the two platforms.

To investigate whether the difference in the quality of sequencing observed in the PCR-free preparations of the *E. coli* genome translated into differences in error rate in sequencing, we analyzed the error rate of the trimmed and filtered mapped reads (AVITI: 3,522,749,978 paired-end reads, NextSeq 550: 2,293,422,615 paired-end reads). Among 571 billion mapped bp sequenced by AVITI, approximately 137.6 million errors were identified. The average error rate of the 17 samples that underwent AVITI sequencing was 0.028%. This represented a more than 89.7% decrease of error rate in comparison with 0.27% from the NextSeq 550 for the corresponding samples (Figure 2C). The full-length perfect read rate of avidity sequencing on the AVITI generated perfect reads ranging from 95.7 to 97% (Supplemental Table 2). A wide variation of perfect reads, however, was found in the samples sequenced on the NextSeq 550 platform (67-96%). However, such large variation in the NextSeq 550 platform disappeared if we included >=148 bp matched: the number of near perfect matched reads ranged from 94.9 to 96.3%, comparable to those from AVITI platform (96.1-98.7%). The wide variation of perfect reads on the NextSeq 550 may relate to the large drop of sequencing quality in the last two cycles of either read. To detect whether the errors from AVITI overlapped with those from NextSeq 550, the position of the errors were plotted across the *E. coli* genome (Figure 2E). As shown in Figure 2E, the concordant errors between AVITI and NextSeq 550 were rare, suggesting that most of these errors were likely generated from the sequencing chemistries rather than the DNA sample preparation since these libraries shared the early stages of sample preparation. Analysis of specific type of nucleotide error showed no preference for a certain type of nucleotide since the errors were evenly distributed among A, G, C, and T (Supplemental Figure 2).

### Short-Read Transcriptome Sequencing

To investigate the utility of AVITI for transcriptome sequencing, we performed short-read RNA sequencing on 9 samples of an external RNA control consortium (ERCC) [12, 13]. ERCC is a standard RNA cocktail of 92 synthetic RNA transcripts ranging from 250 to 2000 nt in length used for quality control analyses. It provides the underlying truth of both RNA quantities as well as the exact sequence of each RNA transcript [14]. ERCC preparations that were sequenced as “spiked-in” libraries on several AVITI sequencing runs generated 32.9 million of filtered mapped paired-end reads, while NextSeq 550 runs where the same ERCC libraries were the primary sample yielded 208.9 million paired-end reads (Supplemental Table 3). As shown in Figure 3A, both AVITI and NextSeq 550, on average, produced excellent correlations with the published quantities of the RNA, with Spearman correlation coefficients of 0.975 for AVITI and 0.982 for NextSeq 550. Of note, the correlation between AVITI and NextSeq 550 was even better: reaching a correlation coefficient of 0.986. The difference of Spearman correlation coefficients between NextSeq 550 and AVITI, however, is not statistically significant (Figure 3B). We then analyzed the quality of the ERCC sequencing by AVITI and NextSeq 550. As shown in Figure 3C and Supplemental Table 1, 88.1% of read 1 and 72% of read 2 from AVITI sequencing reached a per sequence quality score of 40 or better (equivalent to 0.01% error rate or lower). In contrast, no reads from NextSeq 550 were evaluated at this confidence level of accuracy, for reasons previously discussed. While more than 98% of all reads from AVITI reached a per sequence quality score of 30 or better, by comparison, only 90.2% of reads from NextSeq 550 achieved the same values. Similarly, with regards to base sequencing quality, the AVITI platform also showed fewer errors for both base substitution (0.056% versus 0.083%) and insertion/deletion (0.000798% versus 0.002018%) analyses in comparison with NextSeq 550 (Figure 3D). When analyses were performed on error rate per transcript, the results showed that the AVITI had an average error rate of 0.071% per transcript versus an error rate of 0.093% from NextSeq 550 (Figure 3E). When specific transcripts were analyzed, 81 of 92 transcripts showed lower error rates on the AVITI platform vs NextSeq 550 (Figure 3F and Supplemental Table 4).

**Figure 3.**
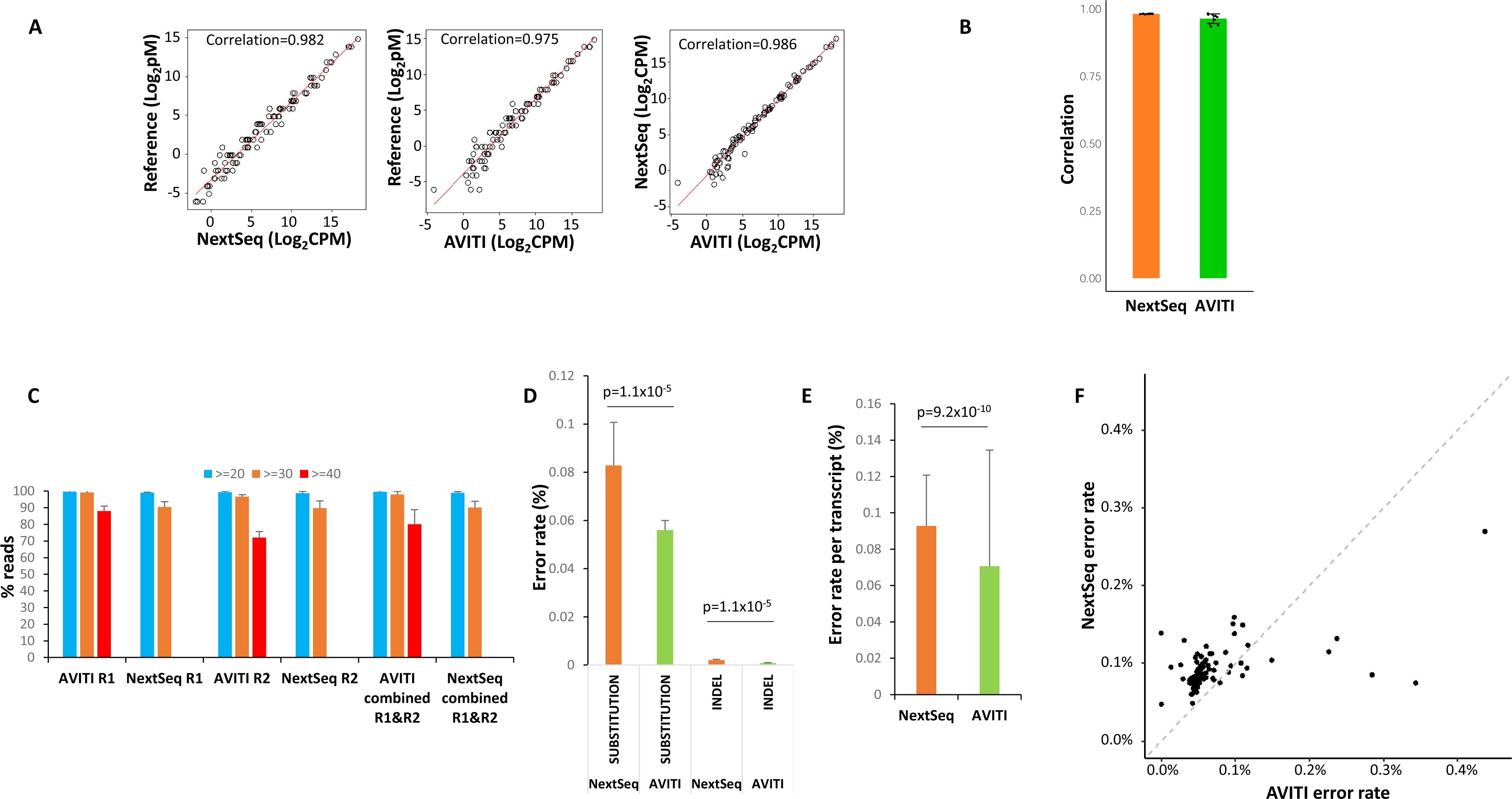
Sequencing of ERCC RNA standard by AVITI and NextSeq 550. (A) Scatter plot of expression quantification of ERCC RNA between reference standard, AVITI quantification and NextSeq 550 quantification. Each opened circle represents the results plotted from the average value of 9 ERCC samples. (B) Correlation of ERCC expression detected by NextSeq 550 or AVITI with standard references. Standard deviations are shown. (C) Per sequence quality score distributions of AVITI and NextSeq 550 sequencing of 9 ERCC samples. Percentage of reads with scores more than 40, 30, and 20 scores are indicated. Standard deviations are shown. (D) Percent error rates of AVITI and NextSeq 550 on 9 ERCC RNA samples. Standard deviations are shown. Wilcox p-values are indicated. (E) Error rates per transcript of AVITI and NextSeq 550 on 9 ERCC RNA samples. Standard deviations are shown. Wilcox p-value is indicated. (F) Scatter plot of error rates of individual ERCC RNA between AVITI and NextSeq 550 sequencing.

### Synthetic Long-Read RNA sequencing

The inclusion of long read sequencing has become more commonplace since its introduction in 2009 [15], and subsequent technological advancements. There are two fundamental approaches to long-read sequencing: native (also referred to as third generation sequencing) and synthetic (SLR). Native long read platforms, such as those from PacBio and Oxford Nanopore, can improve *de novo* assembly, mapping uncertainty and detection of structural variants [16], however, they are also prone to lower per read accuracies, while requiring higher input amounts and yielding less data [17-19]. Synthetic long reads overcome those constraints by harnessing the accuracy and throughput provided by short-read sequencers. Comparisons of LoopSeq synthetic long reads to native long read platforms have previously been performed, and as such are not included in this paper [20-22].

Long-read RNA sequencing is also essential to detect isoform expression and to quantify isoform mutation expression. Previously, we showed that the combination of intramolecular barcoding with short-read sequencing of RNA resulted in highly accurate synthetic long-read sequencing data [21]. To investigate the ability to generate similarly accurate long read sequences on the AVITI platform, ERCC transcripts were intramolecularly barcoded, fragmented, and sequenced. As shown in Figure 4A, long-read sequencing using AVITI and NovaSeq 6000 achieved similar quantification results in relation to the reference standard for all detected transcripts: 0.9617 for AVITI and 0.9588 for NovaSeq 6000. The correlations were slightly lower for both platforms (0.9561 for AVITI and 0.9542 for NovaSeq 6000), if only the full-length transcripts were considered. The base substitution error rate of long-read sequencing from AVITI was 0.051%, while insertion/deletion rate was 0.0075% (Figure 4B). On the other hand, NovaSeq 6000 had a base substitution error rate of 0.054% and insertion/deletion rate of 0.0099%. AVITI also had a slightly lower error rate per transcript than NovaSeq 6000: AVITI had a base substitution error per transcript at 0.054% versus 0.067% for NovaSeq 6000, and insertion/deletion error per transcript at 0.018% versus 0.021% for NovaSeq 6000 (Figure 4C). Fifteen transcripts showed higher error rates on the NextSeq platform while 6 transcripts showed higher error rates in AVITI platform (Figure 4D and Supplemental Table 5). Error position analyses showed higher error rates in the average long transcripts (Figure 4E).

**Figure 4.**
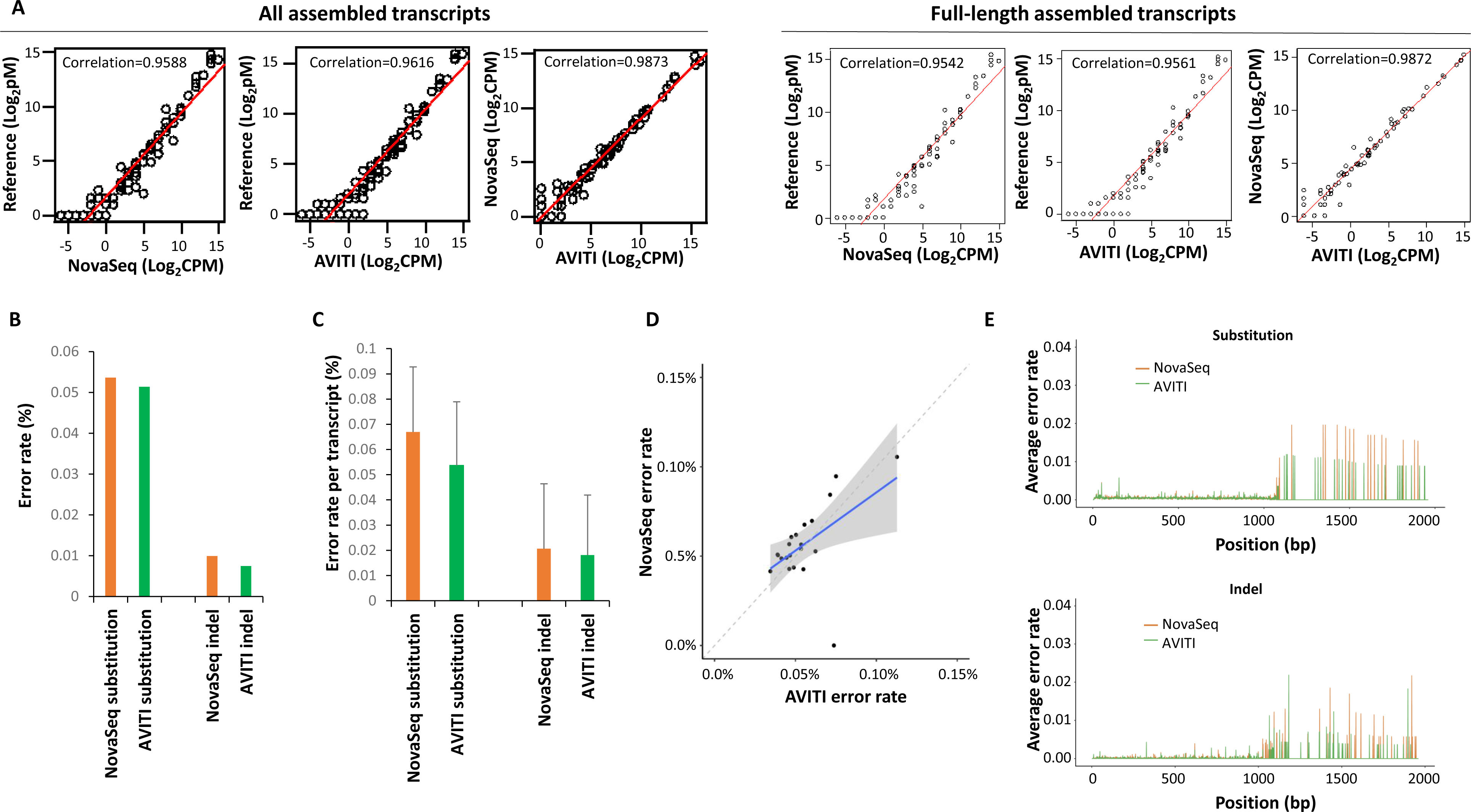
Long-read sequencing of ERCC RNA standard by AVITI and NovaSeq 6000. (A) Scatter plot of expression long-read transcript quantification of ERCC RNA between reference standard, AVITI quantification and NovaSeq 6000 quantification. (B) Percent error rates of AVITI and NovaSeq 6000 on ERCC long-read standard RNA. (C) Percent error rates per long-read transcript of AVITI and NovaSeq 6000 on ERCC standard RNA. (D) Scatter plot of error rates of individual long-read ERCC RNA between AVITI and NovaSeq 6000 sequencing. Blue line indicates linear regression, while the shaded area represents the confidence interval of the linear regression. (E) Distribution of errors by the position of ERCC RNA in AVITI and NovaSeq 6000 sequencing.

### Targeted Single-cell Synthetic Long-Read Sequencing

Long-read single-cell sequencing is an important approach to identify mutation allele expression and isoform switching at individual cell level [23]. To investigate the utility of AVITI with long-read single-cell transcriptome analysis, cDNA samples from a liver cancer patient barcoded with 10x Genomics single-cell 3’ index were ligated to LoopSeq unique molecular index adapters, underwent probe capture, and were subsequently intramolecularly barcoded (Figure 1). These targeted cDNA were then sequenced on the AVITI platform. A more detailed description of the library preparation for this assay is described in Liu et al [24]. As shown in Figure 5A, AVITI sequencing generated an average of 2510 long-read transcripts per benign liver cell and 2426 per hepatocellular carcinoma (HCC) cell (Figure 5A). These were similar to the results generated on an Illumina NovaSeq 6000 platform with the same library preparation: 2418 long transcripts per benign liver cell and 2304 per HCC cell. Similar isoforms and genes per cell were also identified on the AVITI and NovaSeq platform (Figure 5A). The single-cell gene expression correlation between AVITI and NovaSeq platforms were high: Pearson’s correlation coefficients were 0.9549 for benign liver and 0.9776 for HCC (Figure 5B). The cross-platform correlation for isoforms, however, was a bit lower: 0.9192 for benign liver and 0.958 for HCC.

**Figure 5.**
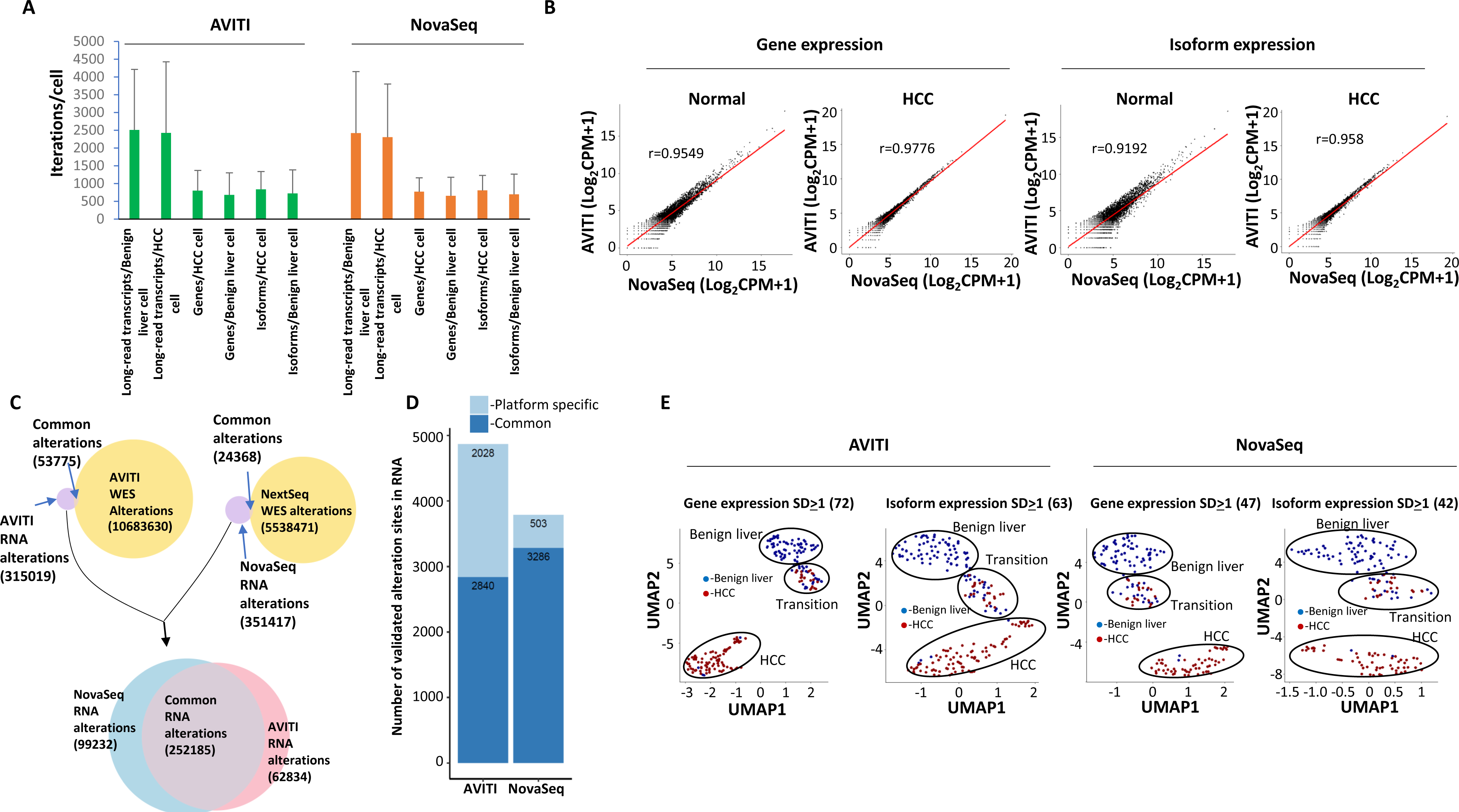
Long-read single-cell sequencing by AVITI and NovaSeq platform. (A) Distribution of targeted long-read single-cell sequencing transcripts, gene, or isoforms per cell by Element AVITI and Illumina NovaSeq platforms. (B) Scatter plots of quantification of long-read transcripts by Element AVITI and Illumina NovaSeq. Pearson correlation coefficients are indicated. (C) Distribution of detected nucleotide polymorphism in long-read transcripts and the genome of HCC by Element AVITI and Illumina NovaSeq. Venn diagram shows the common polymorphisms detected by both genome and transcript analysis. The polymorphisms not detected in the genome of cancer were overlapped to identify sequencing errors specific to each platform. (D) Mutation expression in the long-read single-cell sequencing detected by AVITI and NovaSeq based on the screening of the validated polymorphism against the exome sequencing of gallbladder of the same individual. (E) UMAP clustering of HCC and benign liver cells based on standard deviation of gene or isoform expression across all the cells by Element AVITI and Illumina NovaSeq. Groups of cells of benign liver, transition and HCC are indicated.

Next, we performed single nucleotide polymorphism analysis on long-read transcripts generated by AVITI. As shown in Figure 5C, 53775 out of 368794 single nucleotide polymorphisms (or possible real mutations at the DNA level) expressed in the RNA level were validated by whole exome sequencing of HCC samples on the same platform (Figure 5C). In the long-read transcript data generated using short-reads from a NovaSeq 6000, we were able to validate 24368 out of 375785 SNPs using whole exome sequencing. Interestingly, similarly large numbers of SNPs (AVITI: 315019 and NovaSeq 6000: 351417) present in the RNA level were not present in the cancer genome, suggesting significant RNA editing activity or sequencing errors by the procedure. To investigate which of these non-validated SNPs were platform specific, the non-validated SNPs from AVITI and NovaSeq 6000 were overlapped. The results suggested that 252185 SNPs were common between platforms, indicating their existence preceding the loading of the sequencing cocktail to either platform. These SNPs either represent the results of RNA editing or PCR amplification generated errors. In contrast, 99232 SNPs were NovaSeq 6000 specific while 62834 SNPs were AVITI specific. These platform specific SNPs were generated after the library preparation, and thus, were platform errors. The smaller number of platform-specific errors by AVITI suggests a higher accuracy for the sequencing chemistry. To investigate mutation expression, whole exome sequencing was performed on the genome DNA of gallbladder from the same patient to eliminate the physiological SNP expressions. As shown in Figure 5D, 4868 validated mutations were identified in RNA transcripts using the AVITI, while 3789 mutations were identified with the NovaSeq 6000.

To perform gene-based single-cell analysis, genes with similar expressions across all cells were removed, while genes with expression standard deviations of at least 1 were retained to perform uniform manifold approximation and projection (UMAP) analysis [25, 26]. As shown in Figure 5E, the clustering based on 72 genes from the AVITI platform produced three groups of cells: mostly HCC cells, mostly benign liver cells and a mixture of benign liver and HCC cells (transition). The 47-gene model from Illumina produced similar cell segregation. The 63-isoform UMAP analysis obtained from the AVITI platform produced 3 groups of segregation, albeit a little more dispersed. Interestingly, the 42-isoform model from Illumina platforms produced segregation with larger distance. Overall, both AVITI and the NovaSeq 6000 were comparable in terms of quantifying the transcripts for cell type characterization.

## Discussion

Short-read massively parallel sequencing is fundamental for modern genomics [6]. Prior to 2022, Illumina held the majority of short-read sequencers in the commercial space without an heir apparent [5]. Their platforms utilize sequencing by synthesis (SBS) technology to support numerous applications including whole genome sequencing, RNA-Seq and synthetic long read sequencing. Here, we examine whether the AVITI platform described may present a viable alternative to Illumina sequencing. AVITI’s affinity-based chemistry produces lower requirements for sequencing reagents, higher confidence of base calling, and lower error rate, while maintaining high throughput capacity.

One of the reasons for lower error rates observed with the AVITI platform are the result of separating the sequencing process into two parts: extension of DNA template by synthesis and affinity identification of the nucleotide in the DNA template one at a time, allowing for optimization of nucleotide identification in a separate procedure. Avidite ligands have higher affinity than the nucleotide monomer, resulting in less sequencing reagents employed in the sequencing process. Dissociation constants of the avidite-polymerase-DNA template complex are negligible and minimal background can be achieved, enabling improved imaging without significant decay. These features may enable a large magnitude of improvement in quality over the traditional sequencing methods. In addition, multiple fluorophores of a polymer increase the signal to background ratio may contribute significantly to higher accuracy. Indeed, low error rate in AVITI platform sequencing was also confirmed in human genome sequencing [27]. For a more in-depth description of Avidity sequencing, see Arlsan et al [8].

High quality sequencing usually translates into fewer errors detected in analysis. For short-read *E. coli* sequencing, there was an average of ∼90% decrease of error rate detected in reads from the AVITI platform versus the NextSeq 550. However, such improvement was reduced to a ∼32% reduction when transcriptome sequencing was analyzed. There was an improvement in error rate for the NextSeq 550 platform with regards to transcriptome sequencing which saw a reduction in the error rate from 0.27% in *E. coli* genome sequencing to 0.083% in ERCC transcriptome sequencing, while the AVITI had slight higher error rate in ERCC transcriptome than its *E. coli* genome sequencing: *E.coli* (0.028%) and ERCC sequencing (0.056%). The causes for these variations are unclear. The mild increase of error rate of ERCC sequencing in AVITI platform could be due to errors generated by reverse transcriptase. For synthetic long-read sequencing, the error rate differences between Illumina and Element platforms were statistically insignificant. We speculate that the high coverage per nucleotide may correct most of the errors generated by either platform. Interestingly, most of the sequencing errors between NextSeq and AVITI did not overlap, suggesting a random nature of the errors, rather than specific for certain regions of the DNA sequence.

High quality sequencing translates into high confidence of base-calling and lower errors. Low error rate is important in mutation calling and polymorphism analyses. Due to the comparatively lower confidence of base-calling on current sequencing instruments, the mutation calling threshold has been usually set at a large number of reads. This condition requires an increase in sequencing depth which results in a higher cost of sequencing. In human oncology, mutation calling has formed the backbone of modern diagnostic and therapeutic bases [28, 29]. However, when significant heterogeneity and benign sample dilution are present, ambiguity will be generated. Ambiguity of mutation calling may negatively impact the diagnosis and treatment of cancers. The higher confidence of base calling and lower error rate in avidity sequencing may offer an excellent solution in such situations: a platform with lower error rates may, in the future, help calling correct mutations with fewer reads, and might decrease the cost of sequencing by increasing the signal to noise ratio in mutation detection. It might also increase the number of mutation calls by discovering the mutations previously masked by higher error rates. As a result, avidity sequencing on the AVITI platform may hold promise to become an important role in increasing the fidelity of modern genomic analyses. However, the current study is by no means to demonstrate the superiority of one platform over another since we only selected a few metrics on two organisms and an RNA standard in the analysis. Further study on sequencing of a variety of organisms may be needed to reveal a more comprehensive sequencing capability of AVITI platform.

## Methods

### PCR-Free Whole Genome Short-Read Library Preparation

*Illumina:* PCR-Free libraries were prepared with 1 µg/reaction input of *E. coli* (ATCC 11303) using the KAPA HyperPlus (Cat#: KK8514) reagents with KAPA UDI Adapters (Cat#: KK8726). Instead of utilizing the enzymatic fragmentation reagents provided in the kit, acoustic shearing on a Covaris ME220 platform using 8 microTUBE -50 AFA Fiber H Slit Strip v2 tubes (Cat#: 520240.2) was performed the following settings to produce an average insert size of 350 bp: Sample volume: 55 µL; Waveguide: ME220 Waveguide 8 Place (PN 500526); Temperature Set Point: 12 °C; Repeat/Iterations: 7; Repeat Process Treatment Duration (s): 10; Peak Power (W): 50; Duty Factor (%): 20; Cycles Per Burst: 1000; Avg Power: 10; Total Treatment Time per sample (s): 70. Fifty microliters per reaction of the mechanically sheared gDNA was transferred from the respective AFA tubes into corresponding wells of a clean PCR plate. After adding 10 µl of the prepared master mix, end repair and A-tailing was performed on a Bio-Rad C1000 PCR machine with the following parameters: 30 min at 20°C, 30 min at 65°C. Adapter ligation was conducted by adding 5 µl of the respective KAPA Adapter and 45 µL of the prepared master mix to each well. Reactions were incubated in a PCR machine for 15 min at 20°C with the lid temperature off. Samples were then subjected to a 0.8x SPRI cleanup and a dual SPRI size selection (0.5x/0.66x). All reactions were evaluated for concentration in triplicate using Qubit 1x dsDNA HS assay reagents (Cat#: Q33231) on an Agilent NEO2 Plate Reader and for insert size using HS NGS Reagents (DNF-473-1000) reagents on an Agilent Fragment Analyzer 5300 automated CE platform. Individual reactions of the prepared libraries were normalized for equimolar pooling. The final library pools were evaluated using an Agilent Bioanalyzer 2100 using High Sensitivity DNA Assay reagents (Cat#: 5067-4626). Samples were denatured and diluted for loading on the NextSeq 550 platform per manufacturer’s recommendations.

*AVITI:* The equimolar pooled libraries were concentrated using an Amicon Ultra-30K concentrator column and centrifuging at 14,000 x g for 10 minutes per manufacturer’s guidelines. It should be noted that updates to the Element Biosciences circularization chemistry no longer require the higher input concentration utilized during this preparation. The concentrated library pools were then subjected to circularization with the Element Biosciences Adept Library Compatibility Kit v1.1 (Cat#: 830-00007) according to manufacturer instructions. The circularized library was quantified using qPCR with the provided standard before being sequenced on the AVITI system with 2 x 150 length paired end reads.

### mRNA Transcriptome Short-Read Library Preparation

*Illumina:* For transcriptome sequencing, similar procedures were described previously [30-33]. Briefly, either a 1% dilution of ERCC RNA Spike-In Mix (4456740, ThermoFisher) or fully concentrated samples were processed into a short-read libraries using the TruSeq Stranded mRNA Sample Prep Kit from Illumina, Inc (Cat#: 20020594). The fragmented mRNA was reverse transcribed to cDNA using random primers. This was followed by second stranded cDNA synthesis. The library preparation processes of mono-adenylation, adapter ligation and amplification were performed following the manual provided by the manufacturer. The quantity and quality of the libraries were assessed in a manner similar to that described for whole genome DNA library preparation. The procedure of 2x150 cycle paired-end sequencing on the Illumina NextSeq 550 followed per the manufacturer’s guidelines.

*AVITI:* The linear library pool prepared above (Supplemental Table 3) was amplified with the KAPA HiFi HotStart Library Amplification kit with Primer Mix (#KK2621) to ensure blunt-end products prior to circularization (90s at 98°C; 5 cycles of 30s at 98°C, 30s at 60°C, 30s at 72°C; and 60s at 72°C). Circularization was performed using the Element Biosciences Adept Library Compatibility Kit v1.1 (Cat#: 830-00007) according to manufacturer instructions. The circularized library was quantified using qPCR with the provided standard before being sequenced on the AVITI system with 2 x 150 length paired end reads.

### Synthetic Long-Read mRNA Transcriptome Library Preparation

*Illumina:* LoopSeq preparation of ThermoFisher ERCC RNA Spike-in Mix (Cat#: 4456740) was executed as follows. First, reverse transcription of 11µl ERCC RNA of a 1% diluted template was performed, using a prepared mix of Maxima H Minus RT Buffer and Reverse Transcriptase 200U/µl (Cat#: EP0752, ThermoFisher), RiboLock RNAse inhibitor 40U/µl (Cat#: EO0381, ThermoFisher), 10mM dNTP (Cat#: N0447L, New England Biolabs), LoopSeq specific 100uM Template Switching Oligo (IDT), and a LoopSeq specific 10uM polyTVN UMI-tagging primer oligo (IDT). Incubation conditions were 50°C for 90 min followed by 85°C for 5 min.

UMI diversity in the RT product was then quantified by qPCR. As the RT polyTVN primer oligo includes a UMI sequence, each first strand generated by RT was uniquely UMI-tagged, allowing quantification by qPCR using LoopSeq Amp Mix S and LoopSeq Amp Additive from the Amplicon LoopSeq kit (# 840-10002, Element Biosciences). A master mix of Amp Mix S and Amp Additive was generated following manufacturer instructions and was spiked with 50X SYBR Green (# S7585, Invitrogen) to final concentration of 0.5X SYBR. A dilution series of the RT product was created and quantified by qPCR to determine the dilution needed for an optimized concentration of 15,000 UMIs/µl. PCR was performed for 22 cycles using PCR cycling conditions of 95°C for 3 min, 95°C 30s, 60°C 45s, 72°C 8 min, and a 10°C hold.

After the appropriate dilution was determined by qPCR, twelve replicate 20µl PCR reactions were prepared using LoopSeq Amp Mix S, LoopSeq Amp Additive, and RT product diluted to 15,000 UMIs/µl as template. PCR was performed as described above. 0.6x SPRISelect (Cat#: B23319, Beckman-Coulter) was used for cleanup and generation of a final pool of 20µl. Pooled PCR product was used as input into the Extension LoopSeq kit (Cat#: 840-10003, Element Biosciences).

*AVITI:* Circularization of the synthetic long read transcriptome library prepared above was performed using the Adept Library Compatibility Kit v1.1 (Cat#: 830-00007, Element Biosciences) according to manufacturer instructions. The circularized library was quantified using qPCR with the provided standard before being sequenced on the AVITI system with 2 x 150 length paired end reads.

### Synthetic Long Read Single-Cell Transcriptome Library Preparation

*Illumina:* The Synthetic Long Read Single Cell Transcriptome library was prepared and sequenced as described in Liu et al [24].

*AVITI:* Circularization of the synthetic long read single cell transcriptome library prepared above was performed using the Adept Library Compatibility Kit v1.1 (#830-00007, Element Biosciences) according to manufacturer instructions. The circularized library was quantified using qPCR with the provided standard before being sequenced on the AVITI system with 2 x 150 length paired end reads.

### Bioinformatics Analysis for Short-Read Whole Genome Sequencing Data

*E. coli* DNA sequences were measured by both Illumina and Element Bioscience platforms. For the raw sequencing data, FastQC (https://www.bioinformatics.babraham.ac.uk/projects/fastqc/) [11] was first applied to check the sequencing quality over read cycle. Then, Trimmomatic [34] was employed to trim the adapter sequences and remove the low-quality reads. After the pre-processing, the surviving reads were aligned to the *E. coli* reference genome by Burrows-Wheeler Alignment Tool [35]. BCFtools [36] was then applied to call the variant sites. Sequencing errors (mismatches and small indels) were calculated from the variant sites, excluding the known SNP sites from the *E. coli* strain.

### Bioinformatics Analysis for Short-Read RNA-Seq Data

ERCC spike-in samples were measured by bulk RNA-seq for both platforms. Similar to WGS, the raw RNA-seq data were pre-processed by FastQC [11] and Trimmomatic [34] for quality control and trimming. Then the reads were aligned to the ERCC reference genome by STAR aligner [37]. Transcript counts were collected based on the alignment and the count per million (CPM) was calculated for transcript quantification. BCFtools [36] were applied on the aligned file to call the variant sites. Average error rate was calculated for all the sequencing base-pairs and for each transcript.

### Bioinformatics Analysis for Long-Read RNA-Seq Data

ERCC samples measured by the Loop Seq technology were collected from both platforms. Short-read data produced by each platform were processed using a pipeline to generate LoopSeq synthetic long reads. In this pipeline, adapters were trimmed from the short reads using Trimmomatic [34], short reads were demultiplexed by LoopSeq sample index and UMI, and finally assembled using SPAdes [38]. Synthetic long reads were then aligned to the ERCC reference genome by Minimap2 aligner [39]. The downstream transcript quantification and error rate calculation follow the same pipeline as the ERCC short-read data.

### Bioinformatics Analysis for Targeted Long-Read Single-Cell RNA-Seq Data

Single-cell long-read RNA-seq was performed on a paired tumor – benign sample from an HCC patient by both Illumina and Element Bioscience platforms. Based on short-read sequencing, full length transcripts were assembled into long reads by tool SPAdes [38]. In total, four runs were performed on each library and were pooled together for analysis. Long reads were first annotated by SQANTI [40] based on human genome reference hg38, where each long read was assigned to a known gene and transcript, or novel one otherwise. Then the long reads were demultiplexed by their cell barcode (10X index) and molecule barcodes (LoopSeq index). Eventually, based on the long-read annotation and cell assignment, UMIs were quantified at both gene and isoform level. Valid cells were defined as cells with at least 1000 long-read transcripts per cell. Gene counts per cell and isoform counts per cell were normalized to count per million and compared between the two platforms.

For gene/ isoform expression analysis, R/Bioconductor package Seurat [41] was applied to integrate both benign and tumor libraries. Top genes and isoforms were selected by their expression standard deviation across all the valid cells. Cell clustering was performed based on these selected genes and isoforms, and the cells were visualized by UMAP dimension reduction algorithm [25, 26].

For variant calling, single-cell long reads were aligned to human reference genome hg38 by Minimap2 aligner [39]. Variant calling was performed by BCFtools [36]. For the same sample, whole exome sequencing was measured, and variant calling was performed to detect variant events at DNA level. Variants unique to RNA data are potential RNA editing sites or sequencing errors. Variants detected by both RNA and DNA data are potential mutations.

### Statistical data analysis

All the statistical analyses were performed by R programming. Spearman correlation was calculated comparing pairwise expression levels. The Wilcox test was performed when comparing pairwise error rate.

## Data availability

The raw and processed RNA sequencing data has been deposited to GEO database with accession ID GSE246838 (https://www.ncbi.nlm.nih.gov/geo/query/acc.cgi?acc=GSE246838). The ERCC bulk short-read RNA-seq by Element Biosciences AVITI platform can be downloaded from GSE246830, GSE269800 and Element Biosciences website (https://go.elementbiosciences.com/rna-sequencing-third-party-cloudbreak-freestyle), the ERCC bulk short-read RNA-seq by Illumina NextSeq platform can be downloaded from GSE246832 and GSE269801; the ERCC bulk Loop-seq long-read RNA-seq by Element Biosciences AVITI platform can be downloaded from GSE246834; the ERCC bulk Loop-seq long-read RNA-seq by Illumina NVSQ600 platform can be downloaded from GSE246835; the HCC single-cell Loop-seq long-read RNA-seq by Element Biosciences AVITI platform can be downloaded from GSE246836; The HCC single-cell Loop-seq long-read RNA-seq by Illumina Novaseq platform has been deposited into GEO with accession ID GSE223743 in our previous research. The raw DNA sequencing data has been deposited to SRA BioProject PRJNA1034988 (http://www.ncbi.nlm.nih.gov/bioproject/1034988). The submission includes DNA-seq on *E. coli* by both Element Biosciences AVITI and Illumina NextSeq platforms, and whole exome sequencing on HCC, hepatocyte and gallbladder samples using Element Biosciences AVITI platform.

## Contribution

JHL, SL and TBY conceived the idea. YPY, BGR, TBY, CO, and JHL contributed the experiment design. BGR, CO, JZ, KB, BJK, KM performed the experiments. SL, CO, YPY, JJL, BJK, WW, TBY, and JHL performed the analyses (figures 1-5).

---

**Supplemental Figure 1: Phred score trend line of 16 samples sequenced through AVITI or NextSeq 550.** Phred score from each of the 16 E. coli genome sequencings were plotted cycle by cycle. The average score for each cycle was shown. Sequencing through NextSeq 550 or AVITI was indicated.

**Supplemental Figure 2: Nucleotide error distribution in AVITI and Illumina sequencing.** Left panel: Substitution error in each of A, T, C, and G in NextSeq 550 and AVITI platforms. Right panel: Insertion/deletion error in each of A, T, C, and G in NextSeq 550 and AVITI platforms.

