## Supplemental figures 1-2 for "Utility Analyses of AVITI Sequencing Chemistry"

### Slide 1
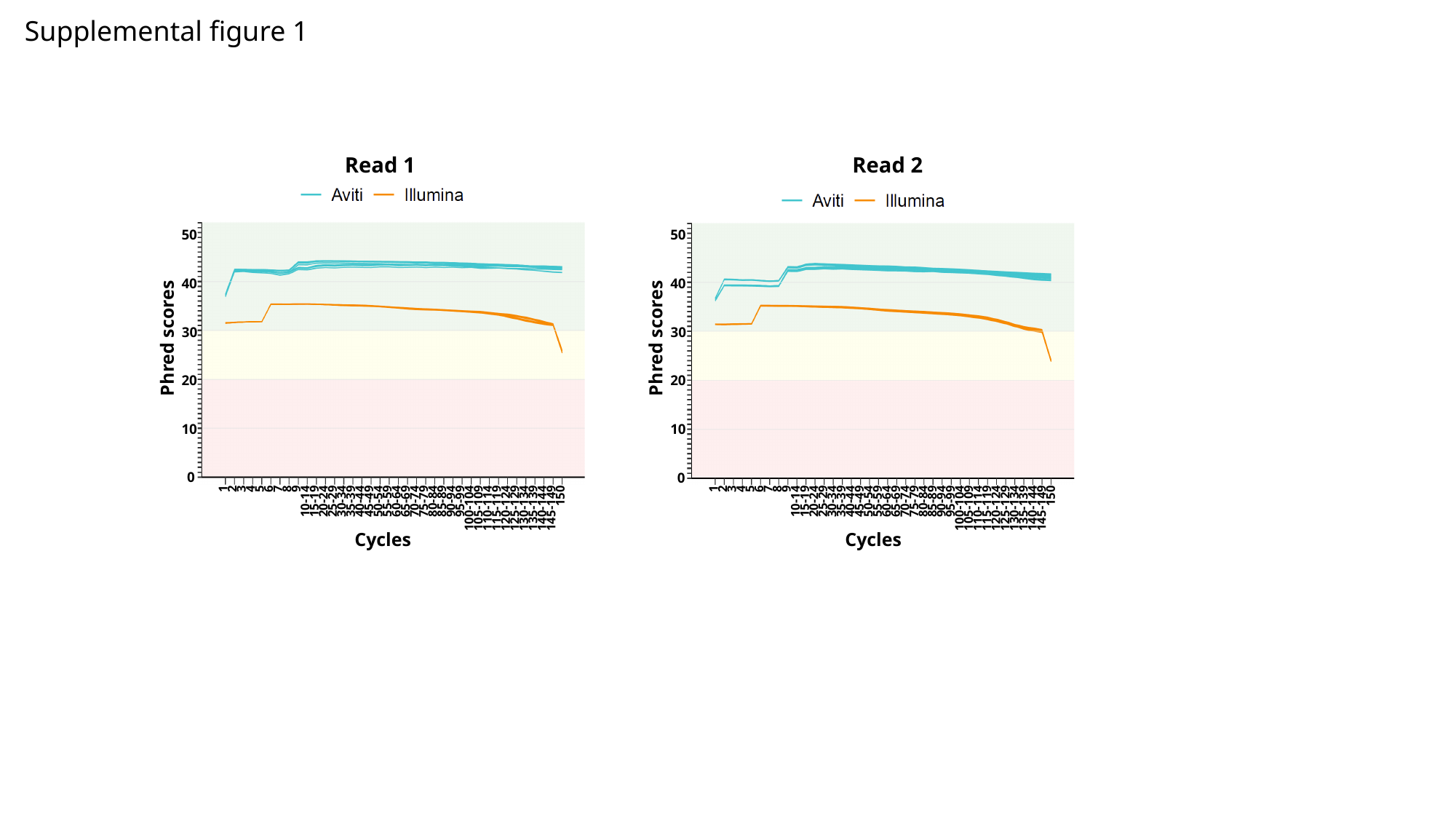

Supplemental figure 1
Read 1			 Read 2
50
50
40
40
30
30
Phred scores
Phred scores
1
2
3
4
5
6
7
8
9
10-14
15-19
20-24
25-29
30-34
35-39
40-44
45-49
50-54
55-59
60-64
65-69
70-74
75-79
80-84
85-89
90-94
95-99
100-104
105-109
110-114
115-119
120-124
125-129
130-134
135-139
140-144
145-149
150
1
2
3
4
5
6
7
8
9
10-14
15-19
20-24
25-29
30-34
35-39
40-44
45-49
50-54
55-59
60-64
65-69
70-74
75-79
80-84
85-89
90-94
95-99
100-104
105-109
110-114
115-119
120-124
125-129
130-134
135-139
140-144
145-149
150
20
20
10
10
0
0
Cycles
Cycles

### Slide 2
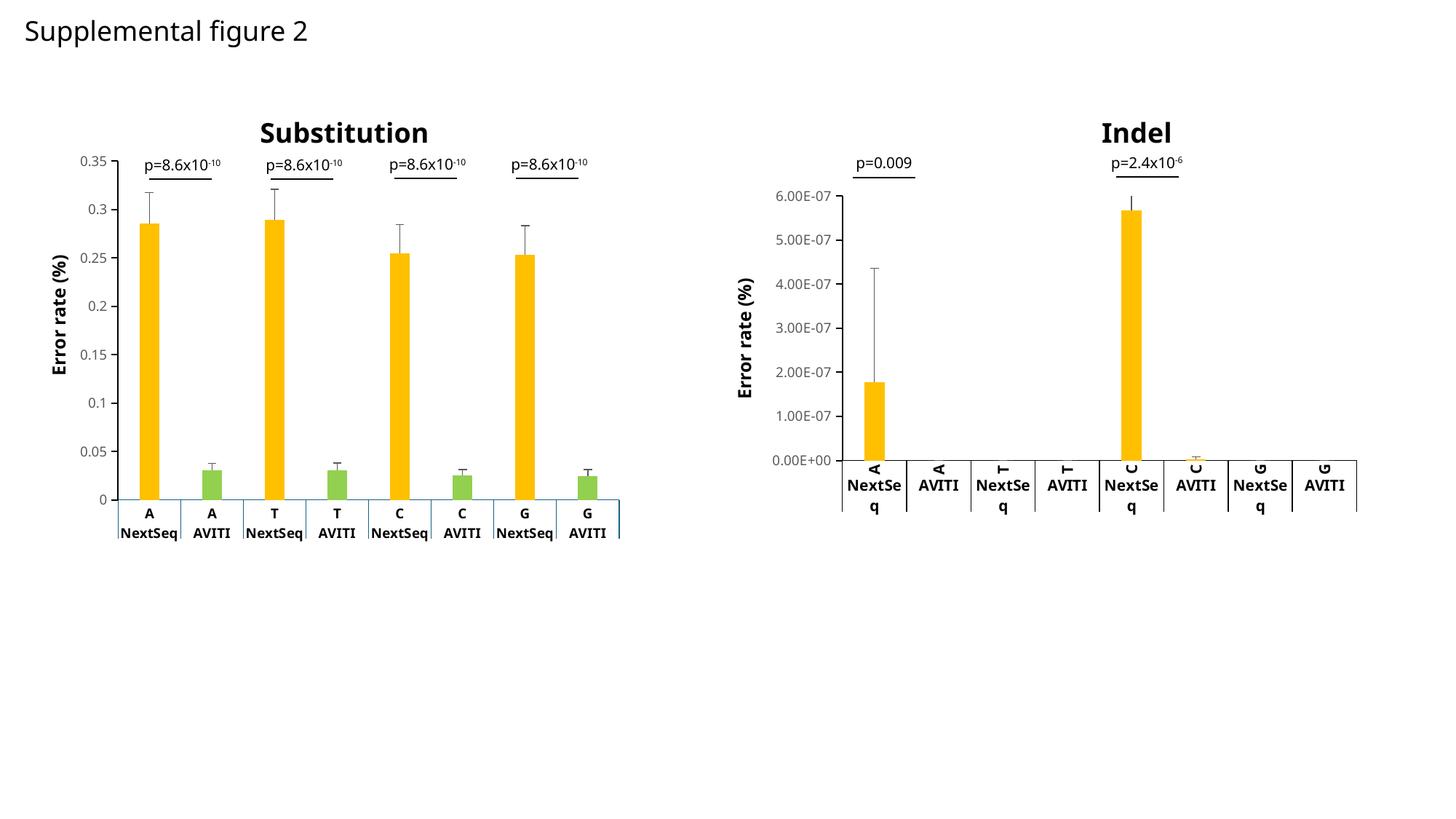

Supplemental figure 2
Substitution					 	 Indel
#### Chart
| Category | |
|---|---|
| A | 0.28552 |
| A | 0.030674 |
| T | 0.288951 |
| T | 0.030840000000000003 |
| C | 0.25468399999999997 |
| C | 0.025127 |
| G | 0.252922 |
| G | 0.024783 |p=2.4x10^-6^
p=0.009
p=8.6x10^-10^
p=8.6x10^-10^
p=8.6x10^-10^
p=8.6x10^-10^
#### Chart
| Category | |
|---|---|
| A | 1.78e-07 |
| A | 0.0 |
| T | 0.0 |
| T | 0.0 |
| C | 5.68e-07 |
| C | 3.07e-09 |
| G | 0.0 |
| G | 0.0 |Error rate (%)
Error rate (%)
